# Facilitation and biodiversity jointly drive mutualistic networks

**DOI:** 10.1101/2020.08.25.266346

**Authors:** Gianalberto Losapio, Elizabeth Norton Hasday, Xavier Espadaler, Christoph Germann, Francisco Javier Ortiz-Sánchez, Adrian Pont, Daniele Sommaggio, Christian Schöb

## Abstract

1. Facilitation by nurse plants increases understorey diversity and supports ecological communities. In turn, biodiversity shapes ecological networks and enhances ecosystem functioning. However, whether and how facilitation and increased biodiversity jointly influence community structure and ecosystem functioning remains unclear.
2. We performed a field experiment disentangling the relative contribution of nurse plants and increasing understorey plant diversity in driving pollination interactions. Both the presence of nurse shrubs as well as increased understorey plant diversity increased pollinator diversity and visitation rates. While nurse and understorey diversity effects on pollinator visitation rates did not interact, the effects of increasing understorey plant diversity on pollinator diversity were stronger in the absence than in the presence of shrubs, meaning that nurse shrubs attenuated the effects of high understorey diversity and buffered the effects of low understorey diversity.
3. We also found positive complementarity effects among understorey species as well as complementarity between nurse plants and understorey species at high diversity. Results also indicate negative selection effects, suggesting that species with generally few pollinators benefit the most in the polyculture, while a species (possibly the nurse plant) with generally lots of pollinators does not. The corresponding changes in pollination networks with the experimental treatments were due to both changes in the frequency of visits and turnover in pollinator community composition.
4. *Synthesis* Plant–plant facilitative systems, where a nurse plant increases understorey plant diversity, are common in stressful environments. Here, we show that these facilitative systems positively influence mutualistic interactions with pollinators via both direct nurse effects and indirect positive effects of increasing plant diversity.
5. Conserving and supporting nurse plant systems is crucial not only for maintaining plant diversity but also for supporting ecosystem functions and services.

## 1 INTRODUCTION

Positive interactions among plants often increase community diversity (Bruno et al., 2003; Callaway, 2007; Levin, 2009; Keddy, 2017; Ellison, 2019). In turn, community diversity increases the functioning and stability of ecological communities (Chapin et al., 2000; Hooper et al., 2005; Tilman et al., 2014; IPBES, 2019) and facilitation contributes to these diversity–ecosystem function relationships (Wright et al. 2017). However, whether and how facilitation and biodiversity interact to influence community structure and ecosystem functioning is not known.

Nurse plants (e.g., leguminous shrubs and trees, cushion plants, kelp) can ameliorate local environmental conditions, increase biodiversity in their understorey and in the entire ecosystem, and organize entire communities (Callaway, 2007; Brooker et al., 2009; Pugnaire, 2010; Butterfield et al., 2013; Cavieres et al., 2014; McIntire & Fajardo, 2014; Rodríguez-Echeverría et al., 2016; Ellison, 2019). By increasing both resources and biodiversity, nurse plants can shape networks of ecological interactions (Losapio et al., 2018) and ultimately influence ecosystem functioning (Losapio et al., 2021). Facilitation has been shown for example to increase plant primary productivity (Wright & Jones, 2004; Michalet & Touzard, 2010; Wright et al. 2017, Schöb et al. 2019) or rates of flower visitation by pollinators (Losapio et al., 2021). In fact, numerous studies have demonstrated the positive effects of diverse flower assemblages on pollination (Laverty & Plowright, 1988; Laverty 1992; O’Connell & Johnston 1998; Feldman et al., 2004; Gazhoul, 2006; Braun & Lortie 2019; Losapio & Schöb 2020). Understanding the influence of nurse species and their positive effects for understorey plant diversity on pollination interactions is crucial for predicting how facilitation can influence ecosystem functioning.

Wright *and colleagues* (2017) proposed three main classes of facilitation mechanisms that positively contribute to ecosystem functioning: (i) indirect biotic facilitation, via reducing species-specific natural enemies (pathogens, herbivores); (ii) abiotic facilitation via nutrient enrichment, such as the legume–rhizobia symbiosis that directly increases nitrogen availability for neighbouring plants; (iii) abiotic amelioration via improving microclimatic conditions. These mechanisms are deduced from literature on biodiversity experiments carried out mainly on plant-species-richness gradients and biomass productivity in temperate meadows (Jiang, Pu & Nemergut, 2008). Thus, the potential of such biodiversity experiments to gather generalizable knowledge is limited to a small and limited range of latitudes, systems, and ecological functions.

Nevertheless, there is an increasing number of studies showing that diversity has similar effects in other types of ecosystems (Albrecht et al. 2012; Fründ et al. 2013; Schleuning et al., 2015; Rohr et al., 2020; Losapio et al., 2021). For example, increasing plant species richness increases pollinator diversity and supports mutualistic network structure (Ebeling et al., 2008; Scherber et al., 2010; Blüthgen & Klein 2011). Analogously, pollinator diversity increases plant fitness and ecosystem services (Albrecht et al., 2012; Fründ et al., 2013; Winfree et al., 2018). As nurse plants increase biodiversity and plant diversity increases pollinator diversity, we hypothesize that the presence of a nurse plant and increased understorey plant diversity, i.e., the two components of a facilitative nurse plant system (e.g., Cavieres et al., 2014; Rodríguez-Echeverría et al., 2016), have joint positive effects on pollinator richness and their flower visitation rates (Fig. 1). Possible mechanisms include microhabitat amelioration by nurse species and polycultures, niche creation and complementarity, cluster and magnet effect, resource sharing, redundancy and diversification of functions.

**Fig. 1.**
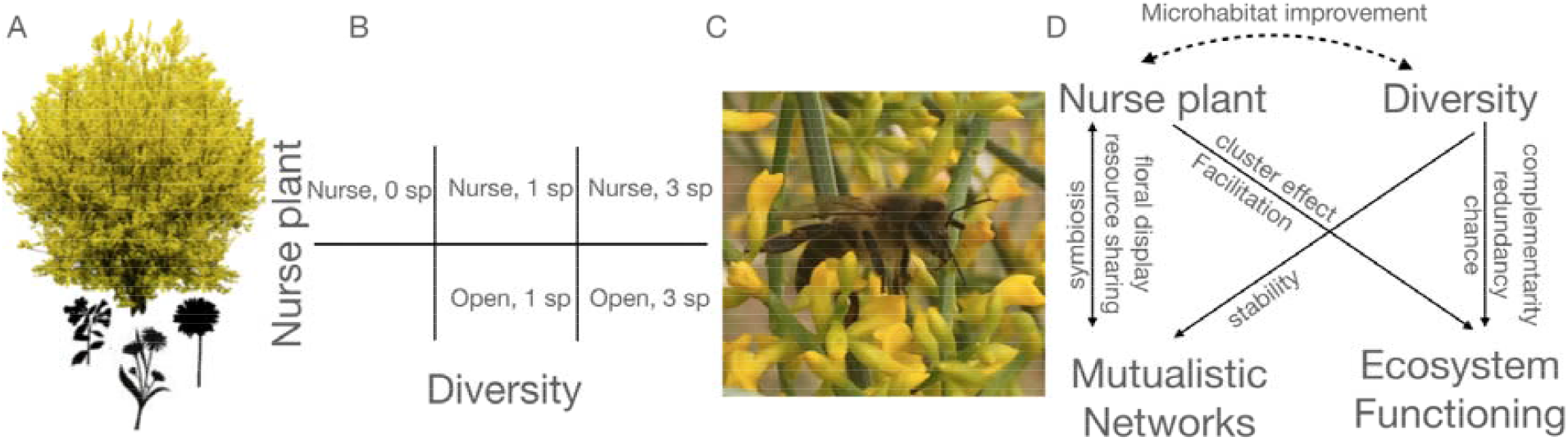
Conceptual framework and experimental design. (A) A legume nurse shrub (*Retama sphaerocarpa*) facilitates diverse herbaceous plant species in the understorey. (B) To examine the joint effects of the nurse plant facilitation and biodiversity change on mutualistic networks, the experimental design includes the treatments of nurse presence/absence and understorey plant diversity. Plant and flower density of understorey species were kept constant. Furthermore, a nurse alone treatment (i.e., shrub without understorey plants) was included. (C) We then examined the response of pollination interactions, such as those between *Apis mellifera* and *Retama sphaerocarpa*. (D) Process pathways among facilitation, diversity, mutualistic networks and ecosystem functioning.

Here, we address this hypothesis and disentangle the relative contribution of nurse plants and understorey plant diversity in driving pollination interactions. We asked the following questions: (i) What are the joint effects of nurse plants and understorey plant diversity on the pollinator community? (ii) What are the costs and benefits for the nurse plant in terms of pollination? (iii) How does a facilitative nurse plant system shape pollination interactions and mutualistic networks?

## 2 MATERIALS AND METHODS

### 2.1 Study system

We used a well-known nurse plant system for the study of plant–plant interactions and particularly facilitation: *Retama sphaerocarpa* (L.) BOISS. (Fabaceae) and its understorey. This leguminous shrub creates “islands of fertility” (Pugnaire et al., 1996; Schlesinger et al., 1996). Mutualistic interactions with *Rhizobia* bacteria in their roots contribute to the improvement of soil resources and amelioration of microhabitat conditions (Moro et al., 1997; Armas, Rodríguez-Echeverría & Pugnaire, 2011). This nurse plant facilitates a wide diversity of understorey species, supports and structures plant communities (Rodríguez-Echeverría et al., 2013, 2016; Lozano et al., 2017; Losapio et al., 2018). Furthermore, *Retama* produces copious yellow blooms pollinated by diverse insects, including small-, medium-, and large-sized Hymenoptera as well as several species of ants (Rodríguez-Riaño et al., 1999).

To experimentally disentangle the direct effect of the nurse plant presence on pollination and its indirect effect through understorey plant diversity, we artificially created gradients of plant diversity in the understorey of *Retama* and in the absence of *Retama*. For this purpose, we selected three annual herbaceous species that commonly grow in *Retama* shrublands at our study site: *Matricaria chamomilla* L. (Asteraceae), *Echium plantagineum* L. (Boraginaceae), and *Carduus bourgeanus* Boiss. & Reut. (Asteraceae). *M. chamomilla* has white and yellow flowers in an open capitulum, *E. plantagineum* has purple tubular flowers along a raceme, and *C. bourgeanus* has blue flowers in a dense capitulum. These three species therefore represent a broad set of flower morphology and pollination niches.

The study was carried out in an oak (*Quercus ilex* L.) savannah in a Mediterranean-type ecosystem at the Aprisco de Las Corchuelas research station in Torrejón el Rubio, Spain (39.81337 N -6.00022 W, 350 m a.s.l., mean rainfall of 637 mm year^-1^ and mean annual temperature of 18 °C).

### 2.2 Experimental design

In order to disentangle the role of direct nurse effects from the indirect effects of the nurse plant on pollinators through increased understorey plant diversity, and to further examine their joint effects on the mutualistic interaction between plants and pollinators, a fully-factorial experimental design was established with two factors (Fig. 1): (i) absence and presence of the legume shrub (open vs nurse), and (ii) understorey plant diversity with zero (0), one (1) and three (3) plant species. This design results in five treatment combinations of open–1, open–3, nurse–0, nurse–1 and nurse–3sp. The treatment combination open–0 was excluded due to the inherent lack of flowers for visitation.

A randomised block design was adopted by grouping together the five treatments and replicating them three times in each block over four blocks, for a total of *n* = 60 plots. Distance between plots within the same block was approx. 1 m. Blocks were distributed randomly over an area of about 4,800 m^2^. Understorey plant species richness was manipulated with plants of the three species transplanted into pots and then the plots placed beneath *Retama* or in the open. The density of understorey plants and flowers was kept constant over all understorey diversity levels. Plants were taken from the neighbouring field and replaced every second day to assure fresh flowers were available. Pots were kept aggregated or separated (*c*. 30 cm apart) below the nurse or in the open. This additional factor (hereafter, aggregation) was replicated per block. In the nurse alone treatment with no understorey species, three empty pots with only soil were placed under the shrub to control for any potential effect of the pots. For the nurse plant treatments, *Retama* shrubs were chosen of approx. the same size (height 127–178 cm and width 125–220 cm). An area of 1 m^2^ at 1 m height was used as the pollinator observation area in each shrub. Flowers of the surrounding vegetation within at least 1 m around each shrub and open area were cleared with a sickle.

Flower visitation was considered as a proxy for the ecosystem function of pollination (Schleuning et al., 2015; Winfree et al., 2018; IPBES, 2019) as it correlates well with seed production (Losapio & Schöb, 2020; but see Lázaro et al., 2013). Flower visits were documented by sampling, identifying, and recording all insects visiting the flowers of each plant in each plot. Observations were conducted between 9 AM and 7 PM over eight days between 1 May and 14 May 2017, covering the blooming phase of the four plant species. Each plot was observed during three surveys of 20 min randomly allocated over the day. Nurse plants and corresponding open plots were observed simultaneously, thereby reducing the potential confounding effects of changing weather conditions within blocks. Each block was sampled within two days. Pollinators were identified at the species level whenever possible, otherwise to the genus. Specimens are conserved in 90% alcohol at our institution collections for inspection and further investigations.

### 2.3 Data analysis

To answer the first question, we calculated the abundance and richness of pollinators in each plot, i.e., at the community level. We assessed the individual and combined effects of facilitation, i.e., nurse presence and understorey plant diversity, on pollinator abundance and richness (two separate models) by means of Zero-inflated Generalized Linear Mixed Modelling (Zi-GLMM) with a negative binomial distribution (Brooks et al., 2017). Understorey aggregation was included as additional factor. Plant species composition and plot nested within block were considered as random effects.

To answer the second question, we calculated flower visits for each single plant species in each plot and conducted cost–benefit analysis. For understorey plants, we assessed the individual and combined effects of nurse shrubs and understorey species richness on pollinator abundance by means of Zi-GLMM with a negative binomial distribution. Understorey aggregation was included as additional factor. Plant species identity and plot nested within block were considered as random effects. For nurse shrubs, we assessed the effects of understorey species richness (nurse alone, 1 species and 3 species; second-degree polynomial) on pollinator abundance by means of Zi-GLMM with a negative binomial distribution. Understorey aggregation was included as additional factor. Understorey composition and plot nested within block were considered as random effects.

To answer the third question, we used a framework based on the variance partitioning of biodiversity effects (Loreau and Hector 2001) for the pollinator community (Losapio et al., 2021). This framework allows comparing the net impact of a diverse plant community on flower visits, distinguishing between complementarity and selection effects (Loreau and Hector 2001). First, we calculated complementarity effects (CE) and selection effects (SE) among understorey species as: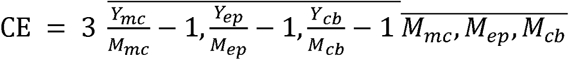 and 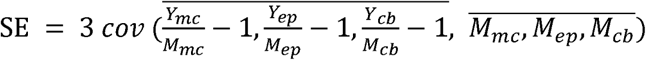, where *Y* and *M* indicate flower visits in polyculture (three understorey species) and monoculture (one of the three understorey species *M. chamomilla* [mc], *E. plantagineum* [ep], *C. bourgeanus* [cb]), respectively. These effects were calculated both in the absence and presence of nurse shrubs. This way, we tested the impact of plant facilitation on CE and SE. Then, the effects of nurse presence, type (CE vs SE) and their interaction on the diversity effect size was tested using a linear model.

Second, we calculated CE and SE between nurse shrubs and understorey species. This way, nurse and understorey were considered as two distinct functional groups. These functional diversity effects were calculated as: 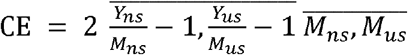 and 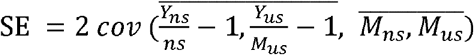, where *Y* and *M* indicate flower visits in polyculture (two functional diversity groups of nurse *ns* and understorey *us*) and monoculture (nurse and understorey alone), respectively. These effects were calculated with both one and three understorey species (nurse–1sp and open–1sp; nurse–3sp and open–3sp). Then, the effect of understorey plant diversity (categorical), effect type (CE vs SE), and their interaction on the diversity effect sizes was tested using a linear model.

Finally, we built mutualistic networks between the four plant species and each of their pollinator species (or genus) according to the additive matrix framework (Losapio et al., 2021). This approach consists of building and comparing observed networks (hereafter, ‘observed’) with ‘additive’ networks. ‘Observed’ networks are built using the plant–pollinator interactions data collected from the empirical plant community, here composed by the nurse shrub and the three understorey species. Instead, ‘additive’ networks are built using data collected from the four treatments of nurse shrub and understorey species monocultures and by pooling plant–pollinator interactions into a single ‘additive’ matrix.

To quantify network structure, we measured network eigenvector centrality (Bonacich, 1987; Csardi & Nepusz, 2006). This metric quantifies the extent to which plant species that are visited by many pollinators are central to the network, or poorly visited plants are strongly connected to few central pollinators. Then, to understand the drivers of differences in mutualistic networks, we measured the dissimilarity between the ‘observed’ and ‘additive’ networks using the framework of beta-diversity of species interactions (Poisot, 2016). In particular, we considered the dissimilarity in species composition and pairwise plant–pollinator interactions. In this case, networks were considered at the block level. Network dissimilarity was calculated within ‘additive’ networks, within ‘observed’ networks, and between ‘additive’ and ‘observed’ networks. Differences among networks were tested in response to the dissimilarity index (species or interactions), dissimilarity within networks nested within dissimilarity between networks, and their interaction using a linear model.

Statistical results are reported in terms of variances explained, using type-II ANOVA (Fox & Weisberg, 2019), and parameter estimates with 95% Confidence Interval. In case of significant statistical interactions, contrasts among factor combinations were computed using estimated marginal means (Lenth, 2020).

## 3 RESULTS

### 3.1 Pollinator community

We found that nurse plants and understorey plant diversity increased the number of pollinators (Fig. 2a) as differences in visitor abundance at the community level were explained by both nurse presence (*P* < 0.001) and understorey plant diversity (*P* = 0.002). Instead, neither aggregation nor the interaction between nurse presence and understorey diversity were significant. *Retama* increased flower visitor abundance on average fivefold compared to open areas (β, 95% CI = 1.72, 0.99–2.45). Furthermore, increasing understorey plant diversity from one to three species increased flower visitor abundance by 82% (0.29, 0.02–0.56).

**Fig. 2.**
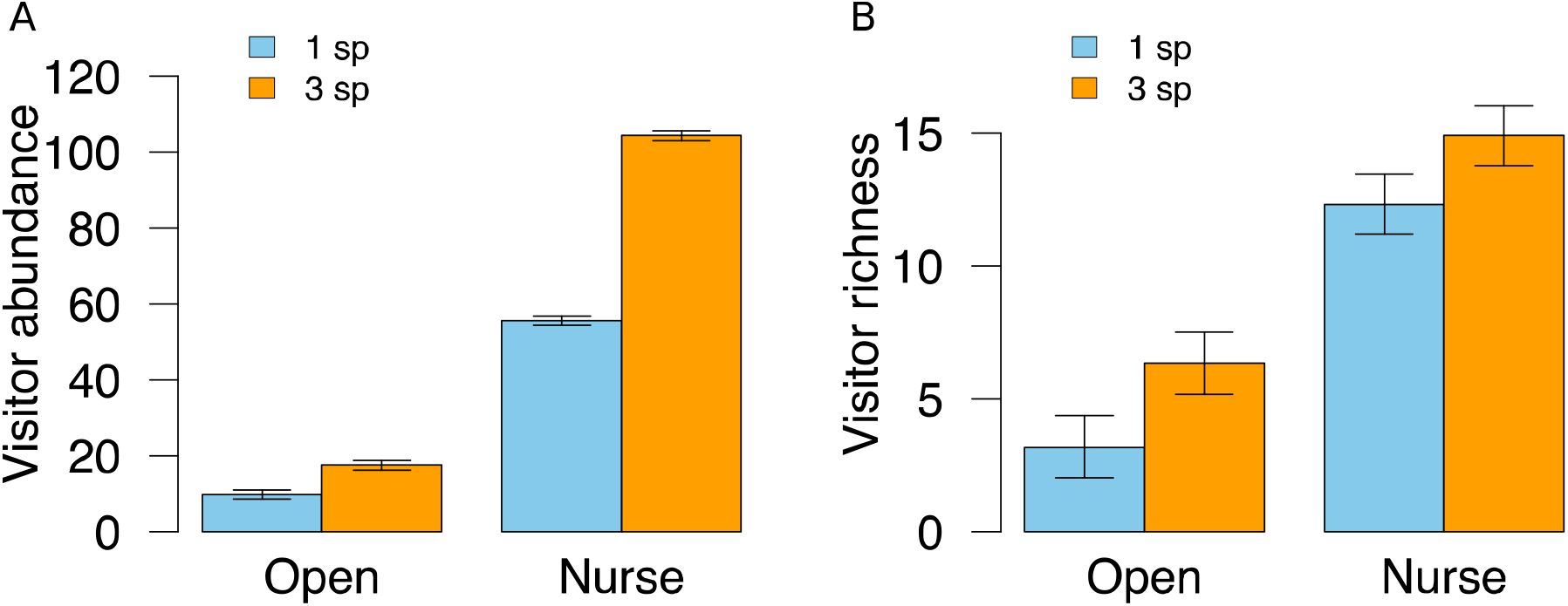
Effects of the nurse plant presence (open vs nurse) and understorey plant biodiversity (1 vs 3 species) on abundance (A) and richness (B) of the pollinator community (i.e., flower visitors) per 20 min per m^2^. Bars indicate standard errors.

Similarly, *Retama* increased the richness of the pollinator community by 74% (1.6, 1.07–2.13, *P* < 0.001), while understorey plant diversity increased pollinator species richness by 24% (0.34, 0.15–0.54, *P* = 0.002). Nurse presence and understorey diversity jointly influenced pollinator species richness (*P* = 0.040, Fig. 2b). The effect of understorey plant diversity was stronger in the open (*c* = 0.69 ± 0.20 SE, *P* = 0.005) than underneath the nurse canopy (*c* = 0.19 ± 0.14 SE, *P* = 0.495), while the effect of nurse presence was stronger at low understorey diversity (*c* = 1.35 ± 0.17, *P* < 0.001) compared with high understorey diversity (*c* = 0.86 ± 0.17 SE, *P* < 0.001). Understorey aggregation had no significant effects.

### 3.2 Benefits and costs

We then explored the effects (i.e., costs and benefits) of nurse plant presence and understorey diversity on flower visitation rate for each species. For understorey species, flowering with nurse shrubs did not have a cost nor a benefit in terms of attracting pollinators (Fig. 3). Instead, understorey diversity had positive effects on visitation rate for understorey plants (*P* < 0.001). These effects were independent of aggregation, nurse presence or their interaction (Fig. 3a). In particular, increasing understorey diversity increased the number of flower visits on each understorey species by 34% on average (0.40, 0.14–0.54). Variance among species was low (0.013).

**Fig. 3.**
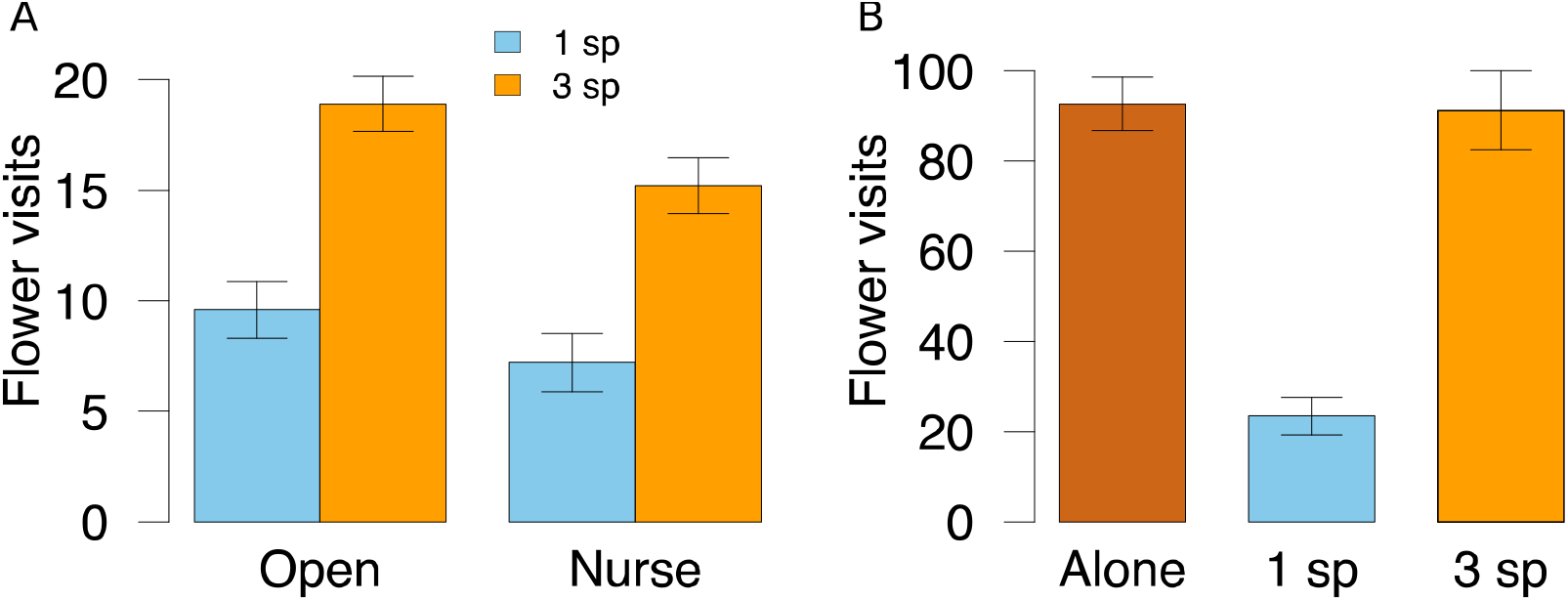
Cost and benefit analysis of nurse plant presence (open vs nurse) and understorey plant diversity (1 vs 3 understorey species) for understorey plants (A) and nurse shrubs (B) in terms of flower visits per 20 min per m^2^. Bars indicate standard errors.

Both aggregation and understorey diversity influenced visitation rate to *Retama* flowers (*P* = 0.045 and *P* = 0.002, respectively). In particular, aggregating understorey plants increased visitor abundance for the nurse shrub by 10% (0.39, 0.01–0.76). Understorey diversity had non-linear effects on flower visitation rate to the nurse plant (Fig. 3b), being positive only at high understorey diversity (quadratic term 2.28, 0.98– 3.59).

### 3.3 Complementarity and selection effects

We then explored diversity effects (i.e., complementarity and selection effects) among understorey species (Fig. 4a) as well as between nurse and understorey plants (Fig. 4b). We found that diversity effects of understorey species were independent of nurse plant presence but varied between complementarity and selection effects (*P* < 0.001). Selection effects were more negative than complementarity effects were positive (Fig. 4a): complementarity effects were marginally positive in the absence and presence of nurse shrubs (499, -33.1–1031; 517, -15.5–1049), respectively, while selection effects were negative in both cases (−754, -1286.3– -222; -771, -1302.9– -238).

**Fig. 4.**
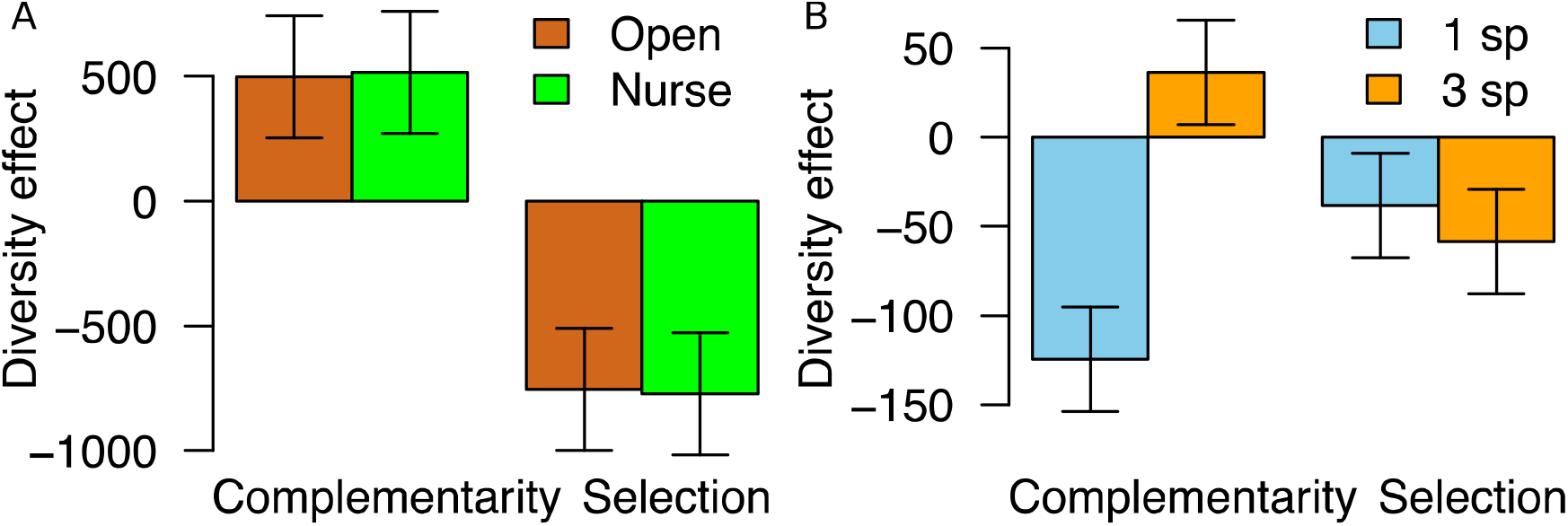
Consequences of nurse plant presence (open vs nurse) for complementarity and selection effects among understorey species (A). Complementarity and selection effects between nurse plants and understorey species (B). Bars indicate standard errors.

Furthermore, diversity effects of combining nurse and understorey plants changed with understorey plant diversity (*P* = 0.033) depending on effect type (*P* = 0.009). While complementarity effects increased with increasing understorey diversity (*c* = 160.0 ± 41.1 SE, *P* = 0.002), selection effects were the same (*c* = 20.1 ± 41.4 SE, *P* = 0.495). Complementarity effects were negative and positive at low and high understorey diversity (−124.6, -188.3– -60.89; 36.2, -27.5–99.93), respectively, while selection effects were on average always negative (−38.5, -102.2–25.26; -58.6, -122.3–5.14).

### 3.4 Network structure

Finally, we explored centrality of the mutualistic network involving plants and pollinators as well as the dissimilarity between additive and observed networks (Appendix S1). We found that observed networks were less centralized than additive networks (β = -0.35, *P* < 0.001, Fig. 5). Looking at network dissimilarity components, we found that species turnover contributed twice as much as interaction rewiring (*c* = 0.20 ± 0.02 SE, *P* < 0.002, Fig. S1).

**Fig. 5.**
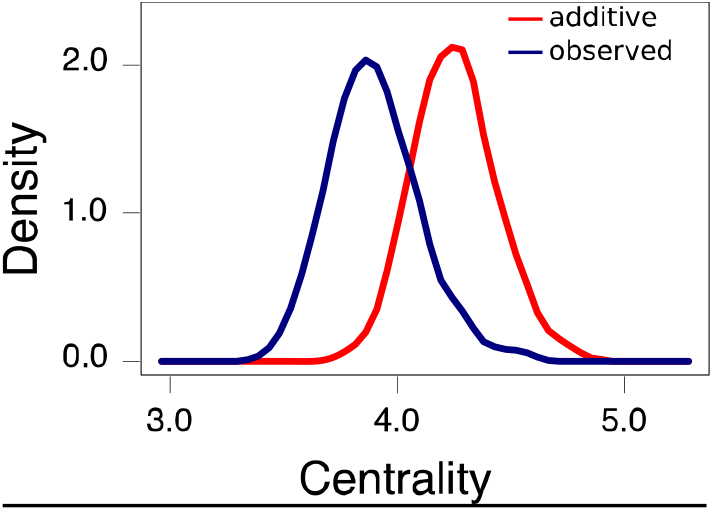
Network centrality of additive networks (red) and observed networks (blue).

## 4 DISCUSSION

Facilitation and species richness play crucial roles in shaping ecological networks and supporting ecosystem functioning, yet their joint effects remain poorly understood. Here, we provide new evidence for the mingled consequences of the presence of a nurse plant and plant diversity for mutualistic networks of pollination, a key ecosystem function. The results of our field experiment confirm the hypothesis that the presence of a leguminous nurse shrub and high understorey plant diversity jointly increased pollinator diversity and visitation rates. Our findings reflect that complementarity and cooperation among different plants resulted in no costs nor benefits in terms of pollination for *Retama* but benefits for pollination of the understorey plant community and the ecosystem overall.

### 4.1 Community-scale benefits

Our results support the hypothesis that nurse plant presence and understorey plant diversity jointly influence pollination, as they both increase pollinator abundance and richness.

Plant facilitation can positively influence ecological networks that include taxa other than plants (Valiente-Banuet et al., 2015), such as pollinators and herbivores (Losapio et al., 2021), ground-dwelling arthropods (van der Zee et al., 2016), mammals (Lortie et al., 2016), and soil microorganisms (Rodríguez-Echeverría et al., 2013; Montesinos-Navarro et al., 2019). Benefits of facilitation involved not only the plant community itself but influenced flower visitors, whose diversity increased in the presence of *Retama* and with increasing understorey richness. Nurse shrubs therefore favour understorey plant diversity, most likely through amelioration of the biophysical environment (Pugnaire 2010), and together with the more diverse understorey plant community also enhanced flower visits via supporting species-rich pollinator communities. Underlying mechanisms for a more diverse pollinator community may be the enhanced floral display via increasing community-level attractiveness to generalist pollinators (Losapio et al. 2021), i.e., ‘cluster effect’ (Krugman, 1991; Laverty, 1992), and resource and service sharing (McIntire & Fajardo, 2014). Moreover, enhanced floral resources via soil symbionts of the leguminous nurse shrub (Harris, 2009; Rodríguez-Echeverría et al., 2016) may also be responsible for improving floral attractiveness in natural conditions.

The fact that increasing understorey plant diversity increases flower visitation rate is consistent with previous biodiversity experiments showing that plant species richness, along with the co-varying factors including blossom cover and presence of particularly attractive flowering species, enhances both the frequency and the temporal stability of pollinator visits (Ebeling et al., 2008; Albrecht et al., 2012). In addition, flower visits increase with plant diversity when diverse flower displays increase the duration of flower provision (Fornoff et al., 2017). Since plant and flower density were kept constant in our experiment, the positive effects of understorey diversity on ecosystem functions are mainly driven by niche complementarity of pollinators and loss of poorly-attractive species, as discussed in the paragraph below (4.2). We used three plant species that have different colours, shapes, volatile cues, nectar and degree of specialization. Such a diversified set of flowers may have contributed to attracting more and diverse flower visitors and have compensated for conditions where the nurse shrub was absent.

Our results show that plant–plant facilitation is an important driver of mutualistic networks. *Retama* increased biodiversity directly and indirectly through facilitation of other plant species, as the increase in plant diversity also increased pollinator diversity, flower visitation rates and ecosystem functioning. The combination of nurse plant presence and associated high understorey plant diversity produced even greater benefits for mutualistic networks than expected by these two factors alone. In fact, the highest levels of pollination were observed in diverse plant communities in the understorey of *Retama*. The presence of nurse shrubs was more important at low diversity, while increasing understorey diversity was more relevant in the absence of *Retama*. Adding more understorey species (at constant plant and flower density) produced stronger effects on pollinator diversity in the absence of nurse shrubs as compared to their presence.

The increased flower visitation rate of communities consisting of *Retama* nurse shrubs and a diverse understorey community compared with communities consisting only of *Retama* or only of herbaceous understorey plants came at no costs for the individual plant species. This shows that *Retama* was not deterring visitors to understorey species. Furthermore, the presence of understorey plants was harmful for *Retama* only at low diversity (one species), while it was neutral at high diversity (three species). These results indicate that diversity decreased the competitive impact of understorey plants on *Retama* visitors, shifting it to non-competitive, neutral effects. A likely process is associated with increasing pollinator attractiveness of a more diverse understorey community following the increase in floral display and flower trait diversity. These results align with studies indicating that the presence of various facilitated plant species can be positive for nurse plant pollination (Laverty, 1992; Braun & Lortie, 2019; Losapio & Schöb, 2020). Increasing understorey diversity to naturally-occurring levels, which are higher than three species, might even shift the effects of understorey plants on *Retama* pollination to positive.

Expanding the understorey diversity gradient would provide further insights into the compensation effect of understorey diversity on pollination of the nurse plant. Beyond that, future studies might want to improve measurements of the ecosystem service by addressing the contribution of flower visits to pollen transfer, visitor effectiveness and seed production. Furthermore, as we controlled for plant diversity and density by using potted plants, our experiment lacks the ‘direct’ facilitation effect of the nurse plant on the understorey plants but allowed us to control for differences in understorey density and floral display in natural communities. We refer here to previous research demonstrating the positive direct effect of *Retama* on understorey plants (e.g., Pugnaire et al., 1996; Rodríguez-Echeverría et al., 2013; Lozano et al., 2017).

### 4.2 Complementarity and selection effects

Our current understanding of the relationships between biodiversity and ecosystem functioning comes primarily from studies focusing on the effects of plant species richness on biomass productivity in temperate meadows (Jiang et al., 2008; Tilman et al., 2014). While competition is often claimed to play a central role in these systems, facilitation is overlooked or lumped within several less explicitly defined processes as complementarity effects (Blüthgen & Klein 2011; Wright et al., 2017). The experimental framework we adopted here (Losapio et al., 2021) allowed us manipulating both taxonomic and functional diversity in combination with facilitation, then measuring complementarity among understorey species as well as between nurse shrubs and understorey plants.

Our results indicate positive complementarity effects among understorey species. Furthermore, complementarity between nurse plants and understorey species increased with diversity. Results also indicate negative selection effects, suggesting that species with generally few pollinators benefit the most in the polyculture (understorey species), while a species (possibly the nurse plant) with generally lots of pollinators does not. These results provide new evidence for a novel facilitation process based on community-scale facilitation (Callaway, 2007; Liancourt & Dolezal 2020) and on the ‘cluster effect’ (Krugman, 1991; Laverty 1992; Losapio et al., 2021): diverse flower assemblages including nurse shrubs are more attractive than monocultures thanks to increased visibility of the community as a whole for attracting a wider spectrum of visitors. This way, being part of the polyculture cluster (nurse shrub with diverse understorey) would increase the chances of being visited, and eventually pollinated.

The positive effects of increased biodiversity for pollination at the community level also translates into increased pollination for each individual species. This is not always the case for biodiversity experiments, where an increase in community productivity does not necessarily follow an increase in species-specific biomass (Tilman et al., 2014).

### 4.3 Network change

By means of the additive matrix framework (Losapio et al., 2021), we compared observed networks that emerge from the plant community as a whole with additive networks that result from pooling plant species as separate components. These two networks appear to be very different, highlighting the non-additivity of nurse and understorey plants. This shows that interactions among plants influence interactions between plants and pollinators. In particular, facilitative interactions among nurse and understorey plants influence pollination networks by changing the identity and frequency of flower visitors. Such nonadditive interactions change network structure, making the network more de-centralised than expected by additive effects, which may ultimately improve the overall resistance and stability of pollinator communities (Blüthgen & Klein 2011). Furthermore, our results show that observed networks were more similar among each other in terms of species interactions as compared to the higher dissimilarity of species interactions observed among additive networks. Drivers of network rewiring are associated with increasing frequency of visits and potentially pollinator population density, thus affecting interaction strength, as well as changes in floral attractiveness and pollination niches, which ultimately promote species turnover and interaction rewiring.

In conclusion, our study shows that plant facilitation systems composed of nurse plants and associated high understorey diversity have joint positive impacts on pollination networks and the functioning of the ecosystem.

## Acknowledgements

GL and CS are grateful for financial support by the Swiss National Science Foundation (P2ZHP3_187938, PZ00P3_148261 and PP00P3_170645).

## Authors’ contributions

GL and CS designed the study; ENH conducted the experiment with the help of GL; XE, CG, FJOS, AP, DS identified the specimens; GL analysed the data and wrote the manuscript with inputs from CS. All authors commented and approved the final publication.

The authors have declared no competing interest.

## Data availability

Data and R code available from the Dryad Digital Repository https://doi.org/10.5061/dryad.bzkh1897m

## Supporting Information

Additional supporting information may be found online in the Supporting Information section.

## Notes

https://datadryad.org/stash/dataset/doi:10.5061%2Fdryad.bzkh1897m

